# Loss of embryonic neural crest cardiomyocytes causes adult hypertrophic cardiomyopathy

**DOI:** 10.1101/270652

**Authors:** Sarah Abdul-Wajid, Bradley L Demarest, H Joseph Yost

## Abstract

Neural crest cells migrate to the embryonic heart and transform into a small number of cardiomyocytes, but their functions in the developing and adult heart are unknown. Here, we map the fates of neural crest derived cardiomyocytes (NC-Cms) and genetically ablate them in embryogenesis in zebrafish. Specific NC-Cm ablation results in aberrant trabeculation patterns and altered Notch signaling, but is not detrimental to the development of the fish or early heart function. Strikingly, embryonic NC-Cm ablation results in adult fish that show severely hypertrabeculated hearts, altered cardiomyocyte size, diminished adult heart capacity and consequently poor physiological response to cardiac stress tests. Thus, we identify a novel developmental mechanism and genetic pathway that predisposes adults to hypertrophic cardiomyopathy and provides the first zebrafish model of adult-onset heart failure.

## Main text

Neural crest (NC) cells are a prototypical stem cell population, migrating from the developing neural tube and capable of transforming into a wide range of cell types during embryogenesis^1^, including cardiomyocytes in zebrafish^2,3^. In humans, syndromes such as diGeorge (22q11 deletion), CHARGE and Noonan/LEOPARD have defects in a variety of NC derived structures, including congenital heart defects (CHDs): Tetralogy of Fallot, Truncus Arteriosus, ventricular septal defects, pulmonary stenosis and hypertrophic cardiomyopathy, implicating NC in human cardiac development^4^. However, it remains unclear whether neural crest-derived cardiomyocytes play a significant role in heart development and disease. The challenge has been to distinguish between primary contributions of NC to cardiac development and sequelae caused by defects in other tissues that subsequently impact cardiac morphogenesis and cardiac function. Distinguishing between global NC versus cardiac NC phenotypes could better inform our understanding of the genetic and developmental etiology of both CHD and adult heart disease. Previous studies have disrupted the cardiac NC population as a whole or different CHD gene candidates within the NC population and then characterized resulting cardiac phenotypes, often in the context of pleiotropic embryonic defects^5–7^. As an alternative and novel approach to decipher NC dependent cardiac phenotypes, we asked how specifically the NC derived cardiomyocyte (NC-Cms) population influences cardiac development and disease, by lineage mapping the NC-Cms and by genetically ablating NC-CMs during embryogenesis. This led us to discover the roles of NC-Cms in regulating the Notch pathway, in patterning trabeculation, and in causing predisposition to adult-onset hypertrophic cardiomyopathy.

### Genetic identification of neural crest derived Cms (NC-CMs)

Several methods have labeled NC before or during migration from the neural tube region and shown that a subset of labeled NC cells are found in the zebrafish heart that are co-labeled with heart-specific markers^2,5,8^. Conversely, we identified NC derived cardiomyocytes (NC-Cms) by genetically marking individual cells that express both neural crest specific genes and cardiomyocyte-specific genes. This dual-component expression both permanently marks the cell lineage and makes it available for temporally-regulated lineage-specific cell ablation (Fig.1A). We generated transgenic lines with a cardiomyocyte-specific driver (*myl7*) of a transcript encoding floxed GFP-Stop followed by tagRFP fused cleavable (P2A peptide) and nitroreductase. Thus, when this transgene is recombined by Cre, it will express red fluorescence, and will allow the expressing cells to be ablated by metronidazole treatment at specific stages of development. This transgenic line was named *Cm:KillSwitch*.

In the absence of Cre-dependent recombination, *Cm:KillSwitch* expresses GFP exclusively in cardiomyocyte lineages (Extended Fig.1). The second transgenic component, called *Tg(sox10:Cre;cryaa:dsRed)*, is a *sox10* driver of Cre expression exclusively in the NC lineages, on a vector marked for selection with cryaa:dsRed for eye expression. We crossed heterozygous *Cm:KillSwitch* adults to heterozygous adults of *Tg(sox10:Cre;cryaa:dsRed)*, selected the offspring that were double-positive for dsRed eyes and GFP hearts (+RE+GFP, Fig.1A). In embryos that carry both transgenes, Cre recombination removes the GFP-Stop and allows expression of tagRFP-P2A-Nitroreductase only in NC-derived cardiomyocytes (NC-CMs), not in other NC lineages or in other non-neural crest derived cardiomyocytes. We found tagRFP+ cells in the heart at 24hpf (hours post fertilization, Fig.1B). These marked cells increased in number until 2dpf (days post fertilization), after which there was no significant increase in the number of NC-Cms (27±3 to 25±2 cells from 2 to 4dpf, n=3 per time-point) or their proportional volume contribution in the developing heart (Fig.1B-I, Extended movie 1). These data indicate that NC-Cm contributes to approximately 10% of the total number of cardiomyocytes in the embryonic 2dpf zebrafish heart^9^. This contribution is early and achieves a steady state by 2 days of embryonic development.

The NC-Cms showed a consistent spatial distribution within the developing heart. NC-Cms localized to the apex of the ventricle, the outer curvature of the atrioventricular canal, and within the border of the inflow tract (Fig.1J-M). In contrast, atrial NC-Cm spatial distribution did not appear as stereotypical as the ventricle. NC-Cm contribution to the inflow tract led us to ask whether the NC-Cms are also integrated into the secondary heart field and whether they contribute to the pacemaker cells of the developing heart^10^. The secondary heart field marker Isl1/2 co-localized with NC-Cms in the proximal ventricle area and a single cell at the inflow tract, suggesting that NC-Cms may also contribute to a subset of pacemaker cells (Extended Fig.2).

### Ablation of NC-Cms

To test the requirement of NC-Cms during heart development, we ablated them during their earliest appearance. Offspring from crosses of *Cm:KillSwitch* and *Tg(Sox10:Cre;cryaa:dsRed)* heterozygous parents were treated with either DMSO (0.5%, control) or 5mM Metronidazole (MTZ) from 30hpf to 48hpf. Only those embryos that were double transgenic, as indicated by dsRed positive eyes and GFP positive hearts (+RE+GFP) were competent to respond to MTZ treatment and ablate the NC-Cms expressing Nitroreductase (Fig.2A). Two controls were included: Sibling embryos that were dsRed-eye negative but GFP positive, treated with MTZ, and double transgenic siblings (+RE+GFP) treated with DMSO. NC-Cm specific cell death was confirmed in +RE+GFP embryos treated with MTZ by immunostaining for activated Caspase-3, a marker of cell death. No significant cell death was observed in the two control groups (Extended Fig.3).

After treatment and subsequent washing at 48hpf, embryos were grown to 5dpf and hearts were analyzed by confocal microscopy. Importantly, no new tagRFP+ cardiomyocytes were observed three days after the ablation period, but an occasional extruding remnant of a tagRFP+ NC-Cm was detected (Fig.2B-D). These results indicate efficient ablation of the initial NC-CM population by 48hpf, and that no new *sox10*-expressing cardiomyocytes were produced, either by subsequent waves of NC migration or by *de novo* expression of *sox10* in other cardiomyocyte lineages. While a previous study implicated two waves of cardiac NC migration to the heart, an early wave and a late (post 3dpf) wave^5^, our data suggest that any late waves of NC migration into the heart do not contribute to cardiomyocytes and/or do not express the NC marker *sox10*. Persistent (past 3dpf) *sox10* expression has only been reported in peripheral glia cells in zebrafish^11^, thus it is more likely that any later wave of *sox10* positive, NC derived cells do not transform into myl7-expressing cardiomyocytes.

### NC-Cm ablated hearts have aberrant trabeculation

Given that approximately 10% of the total number of early cardiomyocytes are NC-Cms, it was surprising that no gross morphological phenotypes were observed in the NC-Cm ablated embryonic hearts (+RE+GFP, MTZ treated) compared to control-treated hearts (Fig.2B-D). We found no significant differences in ventricle size or heart rate in NC-Cm-ablated embryos at 6dpf and juveniles at 14dpf (Extended Fig.4), well beyond the age at which mutants with severe heart defects can survive^12^. However, ventricles of the NC-Cm deficient hearts had a subtle internal defect compared to controls. Analysis of confocal slices of the ventricles from NC-Cm ablated hearts at 5dpf demonstrated an unusual disarray of trabeculation compared to control hearts (Fig.2E). We quantified this phenotype by measuring the angle of the primary branch of the trabeculae as they contact the inner ventricular surface proximal to the atrioventricular canal (Fig.2E bottom panel, Extended Fig.5). The position of each trabecula was measured relative to the anterior-posterior coordinate position (0 to 1.0) in the ventricle (Fig.2E). A significant difference was found in the trabeculation pattern of the NC-Cm ablated hearts compared to both sets of controls (Fig.2F).

### NC-Cm relation to Notch regulation of trabeculation

Notch signaling is an important regulator of trabeculation during heart development. A recent report demonstrates that Notch activated cardiomyocytes signal to their immediate neighbors to trigger the initiation of trabeculation in the neighboring Notch-negative cardiomyocyte^13^. Given the aberrant organization of trabeculation patterns in NC-Cm-ablated hearts, we explored the spatial relationship of Notch signaling and NC-Cms by crossing a *sox10* reporter line *Tg(sox10:tagRFP)* with the Notch signaling reporter line *Tg(TP1:d2GFP)* and examined the resultant embryonic hearts by confocal microscopy^14^. Transiently tagRFP labelled NC-Cms were not co-incident with Notch activated cells; cells were not found to co-express both reporters (Fig.3A). Instead, NC-Cms in the ventricle were immediately adjacent to Notch activated cardiomyocytes (Fig.3A, white arrows). This result suggests that the NC-Cms could be the Notch ligand providing cells, which trabeculate, as described in the previous model, and as illustrated in Figure 1D. Therefore, we asked whether Jag2B, which is thought to signal to Notch activated cells, and Erbb2, which is involved in repression of trabeculation ^13,15^, are disrupted in NC-Cm ablated hearts.

Isolated, embryonic 4dpf hearts from NC-Cm ablated fish and their sibling controls were used for quantitative PCR analysis of *erbb2* and *jag2b*. *Jag2B* was downregulated in NC-Cm ablated hearts while *myl7* and *nrg2a* expression were not significantly changed compared to treated, control hearts (Figure 3B). *Erbb2* expression was not significantly affected in NC-Cm ablated hearts compared to control hearts, perhaps because the expected increase in *erbb2* expression, upon NC-Cm and Notch signaling disruption, is counteracted by the loss of *erbb2* expression from the NC-Cms themselves in NC-Cm ablated hearts^16^.. Together, these data support and extend the current Notch/Neuregulin model of cardiac trabeculation, and lead us to propose that the NC-Cms are critical for the correct patterning of ventricular trabeculation by providing a unique cellular source of Jag2B (Figure 3C).

#### Embryonic ablation of NC-Cms predisposes adults to heart failure

Ablation of NC-Cms resulted in a subtle but consistent trabeculation defect in embryos and juveniles but did not diminish viability, and the embryos grew to adulthood (n=16/20 for MTZ treated −RE+GFP controls compared to n=20/22 for MTZ treated +RE+GFP siblings). The effects of altered embryonic trabeculation on adult cardiac structure and physiology have not been reported. We grew the NC-Cm ablated embryos and sibling controls to adulthood and analyzed their hearts by whole mount fluorescence imaging. Control hearts had large patches of tagRFP+ NC-Cms in the apex of the heart, indicating that NC-CM lineages persisted into adulthood, with similar topological distribution (Fig.4A), and confirmed by flow cytometry (Extended Fig.6). The embryonically ablated NC-Cm hearts (+RE+GFP, MTZ treated), had negligible tagRFP fluorescence in the heart, indicating that no subsequent (post 2dpf) contributions of NC cell lineages persisted in the heart into adulthood, and that *de novo sox10* expression did not occur after the initial embryonic NC-Cm contribution (Fig.4C; confirmed by FACs Extended Fig.6). Remarkably, sections of the NC-Cm-ablated hearts revealed a massive hypertrabeculation (Fig.4D-F), more substantial than that predicted by the subtle alterations in the patterning of embryonic trabeculation. Trabeculation was quantified by determining the percentage of area in a ventricle section covered in trabeculae, which was ~70% in control hearts and ~85% in NC-Cm ablated hearts (Fig. 4G). This measurement is inversely correlated with the luminal area that is available for blood flow through the ventricle, which was decreased two-fold, from 30% in controls to 15% in NC-Cm ablated hearts.

This hypertrabeculation phenotype is analogous to hypertrophic cardiomyopathy in humans, which can be due to either increased cardiomyocyte size, increased cardiomyocyte cell number or both^17,18^. We therefore quantified cardiomyocyte number and size in NC-Cm ablated adult hearts compared to control siblings. Using Mef2 antibody staining along with DAPI to specifically demarcate cardiomyocyte nuclei in ventricle sections of the adult hearts, we found no significant difference in the number of cardiomyocytes per trabeculae area in NC-Cm ablated hearts compared to their control siblings (Figure 4H-J, Extended Fig7A). To determine if cell size was altered, ventricles from treated and control adults were isolated, dissociated into single cells and cultured in chamber slides for 24 hours to allow adherence of dissociated single cardiomyocytes. The cells were then fixed and stained for GFP and DAPI and GFP positive cardiomyocytes were imaged and analyzed for cell area. Cardiomyocytes from NC-Cm ablated ventricles were significantly larger in area than their sibling control cardiomyocytes and this increase in area was most likely because of their significant increase in overall cell length (Extended Fig.7B-D, n ≥ 25 cells per condition, p=0.02 for area and p=0.03 for length measurements). This result, increase in cardiomyocyte size versus number in NC-Cm ablated hearts, was also confirmed by flow cytometry analysis of dissociated ventricles from NC-Cm ablated adults and sibling controls (Extended Fig.8). Together, these results indicate that increased cardiomyocyte size contributes to the hypertrabeculation/ hypertrophic cardiomyopathy phenotype in adults raised from NC-Cm ablated embryos.

Because the NC-Cm ablated hearts were dramatically hypertrabeculated, with diminished lumen volume for blood flow, we asked whether embryonic ablation of NC-Cms had an impact on adult cardiac function. NC-Cm ablated adults and sibling controls were subjected to a cardiac stress test via a swim tunnel assay, in which adult fish are challenged with step-wise increases in water speed, analogous to step-wise increases in treadmill speed in stress-tests of human adult cardiology patients. The swim tunnel assay measures the critical water speed at which individual fish fatigue in swimming (Extended Movie 2). NC-Cm ablated adults performed significantly poorer in the swim tunnel assay than their sibling controls and fatigued at much lower water speeds (Fig.4K, 4L). These results indicate that early and subtle defects in embryonic heart development, dependent on NC-Cm lineages, can predispose adults to performance-induced heart failure.

Overall these findings demonstrate previously unknown roles for NC derived cardiomyocytes, using unique lineage labeling and genetic ablation approaches. Importantly, the hypertrophic cardiomyopathy and heart failure in adults and aberrant trabeculation patterning in embryos can only be attributed to the post-migratory NC-derived cardiomyocytes in our study. This is in striking contrast to previous studies that arbitrarily ablated embryonic ventricular cardiomyocytes and reported no consequential effects on subsequent embryonic heart regeneration, function and trabeculation formation^19,20^. Our study demonstrates that the NC-Cms are not a replaceable cardiomyocyte population, unlike other cardiomyocytes of the heart, and supply a required, innate function of regulating trabeculation in the ventricle in the embryo and cardiac structure and function in adulthood. An additional observation from our results is that while the NC-Cm function in trabeculation is required, the ventricle is still capable of maintaining its’ steady state population of cardiomyocytes, as we found no significant difference in cardiomyocyte quantity despite the ~10% loss of cardiomyocytes by NC-Cm ablation. Previous attempts to analyze the consequences of NC ablations reported changes in embryonic ventricle morphology, heart rate and other defects that we did not observe^3,5,21^. Those reported effects were likely due to secondary effects of perturbation of other NC-derived lineages that contribute to other embryonic structures such as aortic arches or endocardium, resulting in pleiotropic phenotypes that can have secondary effects on heart function^5,8,21^.

Our findings extend and clarify the roles of Notch/Neuregulin regulated trabeculation in the ventricle during embryonic development. Previous models did not suggest mechanisms by which certain cardiomyocytes were pre-patterned to express Notch signals, which then subsequently trigger neighboring cells to Notch activation and suppression of trabeculation^13^. From our observations of NC-Cm lineage distribution in the ventricle in relation to Notch-responding cardiomyocytes, and the results of altered Notch ligand expression and altered patterning of trabecula in NC-Cm ablated hearts, we suggest that NC-Cms serve as a pre-specified source of Jag2B presentation in the 3dpf ventricle, which then impacts the spatial patterning of trabeculation. Thus, in the absence of NC-Cms, normally patterned presentation of Jag2B is diminished and trabeculae are disorganized. While NC-Cms are not known to comprise a significant portion of the mammalian cardiomyocyte population^22–24^, their roles in the patterning of trabeculation and in adult heart function have not been explored. In humans, the etiologies of hypertrophic cardiomyopathy and heart failure are unknown^17^, and our study proposes a novel neural crest derived cardiomyocyte (NC-CM) component. Our results also suggest that the hypertrophic cardiomyopathy phenotype prevalent in the NC disease, Noonan’ syndrome, could be a direct a consequence of perturbation of the NC-Cm population^25,26^. If NC function is more broadly affected than just NC-Cms, other non-cardiac phenotypes will also present, such as in the range of neurocristopathies that are clinically evident. However, if just NC-CMs or the genetic regulatory pathways expressed therein are more specifically altered, either by mutation or by developmental defects, individuals could have subclinical defects that only become apparent as stress-induced adult-onset heart failure. Further studies of the genetics and developmental regulatory mechanisms of NC-Cms will inform our understanding of the various presentations of CHDs, their relation to and their roles in adult cardiac function.

**Figure 1.**
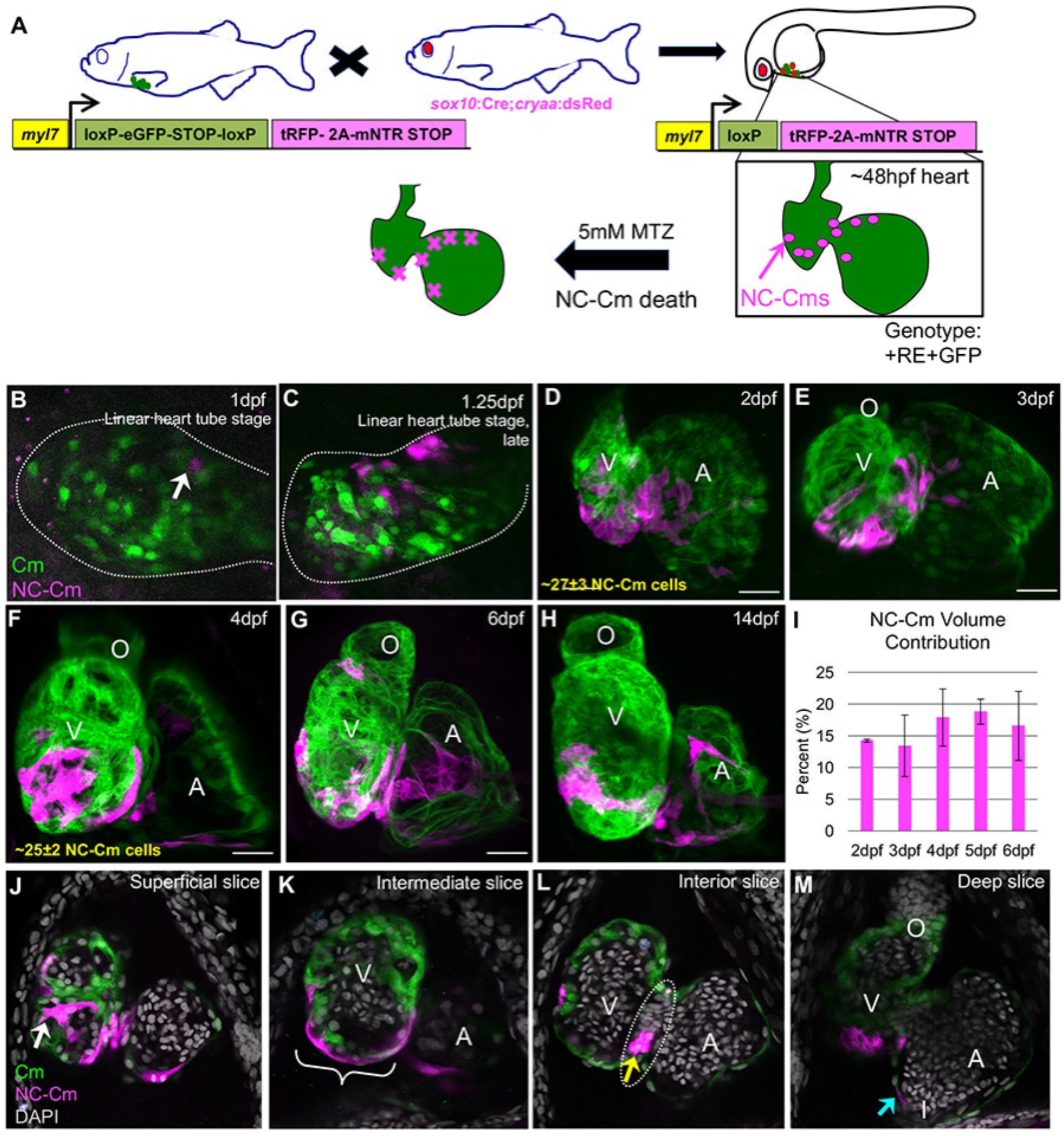
NC-Cm during heart development. A) Schematic of the NC-Cm lineage labeling and ablation setup using the *Cm:KillSwitch* transgenic (myl7-driven transgene) crossed to the NC driver *Tg(sox10:cre;cryaa:dsRed)*. Metronidazole (MTZ) treatment causes mNTR-expressing cells to die, i.e. NC-Cms switched to express tagRFP+ and mNTR. B-H) Contribution of NC-Cms to the developing heart over time. Confocal maximum intensity images of each development stage (1-14dpf). On average, 27±3 NC-Cms were found at 2dpf and this number did not significantly increase by 4dpf (25±2 NC-Cms). Quantification was from confocal 3D stack images at indicated timepoints and from three individuals. Dotted line outlines heart tube. I) Graph displays average percent of total cardiomyocyte volume contributed by tagRFP cardiomyocytes (NC-Cms) at indicated developmental times; three or more individual hearts at each time point. Error bars are standard deviation. J-M) Confocal slices of a 4dpf heart from NC-Cm lineage labelled embryos. White arrow indicates trabeculating NC-Cm. Bracket denotes the apex of the ventricle. Dashed line indicates the AV canal and the yellow arrow indicates the couple of NC-Cms found on the outer curvature of the AV canal. Blue arrow indicates the NC-Cm found at the border of the inflow tract. O=outflow tract, V=ventricle, A= atrium. Scale bar is 30uM. Images are representative of n≥3.

**Figure 2.**
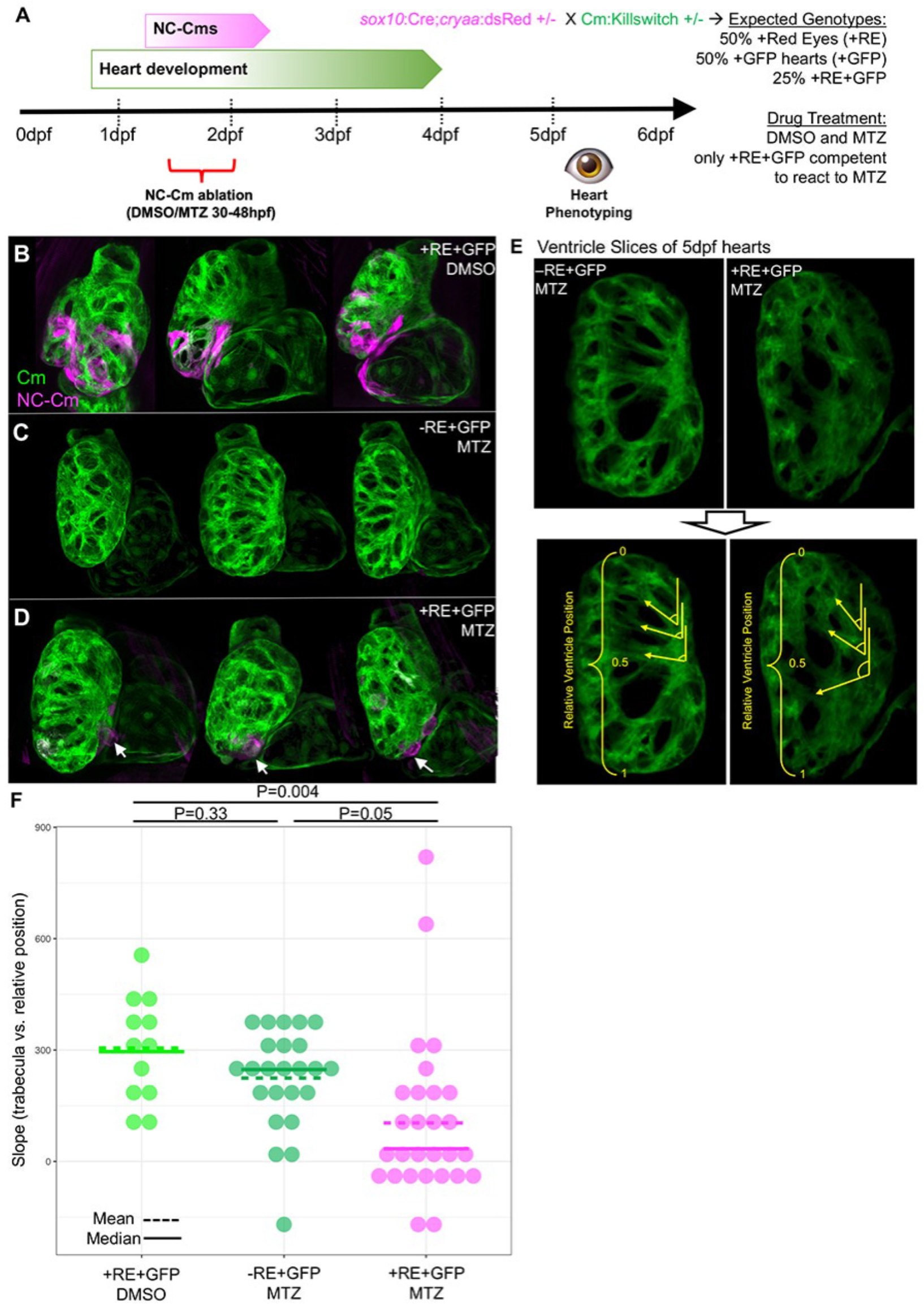
NC-Cm ablation and heart development. A) Schematic of NC-Cm ablation protocol. Transgenic, heterozygous adults for *Cm:KillSwitch* and *Tg(sox10:cre;cryaa:dsRed)* were crossed to generate a pool of siblings with three transgenic genotypes: only *Cm:KillSwitch* positive (+GFP); *Tg(sox10:cre:cryaa:dsRed*) positive (+RE); or both (+RE+GFP). Embryos that were positive for both transgenes were treated with either DMSO (control) or MTZ from 30-48hpf. Only this double-trangenic genotype was competent to respond to NC-Cm ablation by MTZ. Sibling embryos that were only positive for *Cm:KillSwitch* (−RE+GFP) were also treated with MTZ as a drug control. After treatment, embryos were phenotyped at 5dpf. B-D) Results of control and NC-Cm ablation in 5dpf hearts. Confocal maximum intensity projection images from three hearts from each condition. NC-Cm detection by tagRFP fluorescence was largely absent from the MTZ treated +RE+GFP embryos compared to their DMSO treated sibling controls (D compared to B). White arrows indicate a remnant, extruding NC-Cm as a consequence of cell death. E) Trabeculation analysis of NC-Cm ablated hearts. Confocal slices of embryos subject to NC-Cm ablation as in A) were analyzed for trabeculation patterns. Control hearts had an array of trabeculae with primary branches arranged along anterior-posterior coordinate. In contrast, NC-Cm ablated ventricles had poorly organized trabeculae (E upper panel left compared to right). The angle of the primary branch of a trabecula and relative anterior-posterior position of the primary branch within the ventricle were measured (E bottom panel, yellow arrow lines depict a primary trabecula branch and relative position in ventricle is represented by yellow bracket). The position and angle of the primary trabeculae branches were measured relative to the AV canal. These data were collected for controls and NC-Cm ablated hearts (+RE+GFP, MTZ treated) and a slope was computed using the trabecula angle to position data for each individual heart (see Extended Figure 5). F) Computed slope values for individual hearts in each condition. Mean is indicated by the dashed line and median by the solid line. The slope measurement was significantly different for NC-Cm ablated hearts compared to their sibling controls. P-values computed by TukeyHSD on ANOVA(F(62,60)=3.31).

**Figure 3.**
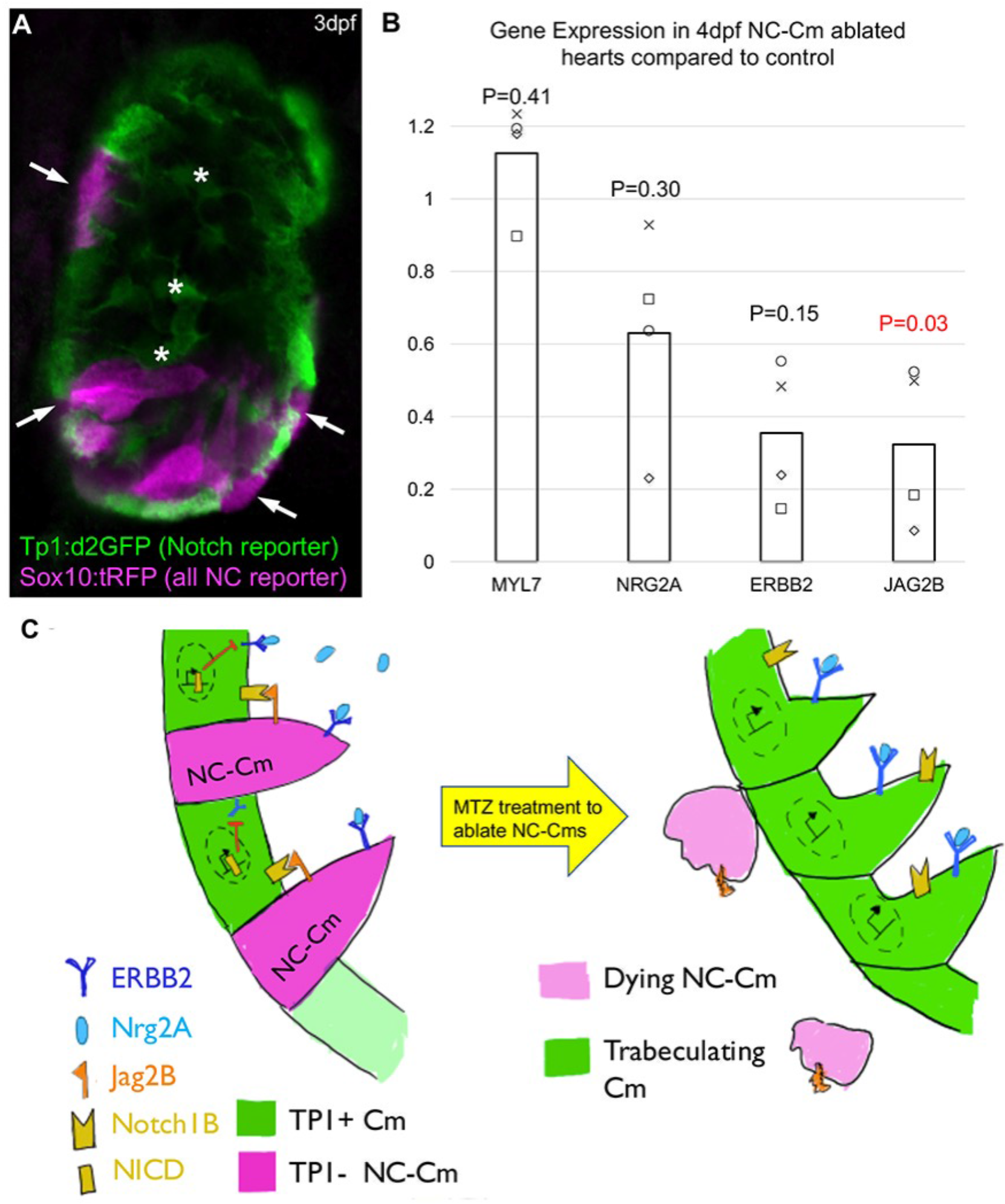
NC-Cm cells regulate Notch signaling during trabeculation. A) Ventricle section of a 3dpf embryo from a *sox10:tagRFP* transgenic line crossed with the Notch reporter Tp1:d2GFP line. NC-derived lineages (tagRFP) did not show high levels of Notch response (d2GFP). Arrows indicate trabeculating, sox10+ cardiomyocytes i.e. NC-Cms that were next to a Notch activated cardiomyocyte. Internal, rounded GFP+ cells are Notch activated endocardial cells (white asterisk). B) qPCR gene expression of isolated 4dpf hearts from NC-Cm ablated and control siblings. Values are delta delta Ct computations. Bars represent average of four biological replicates. Points are individual experiments. Delta Ct values were normalized to Rpl11 expression and used to compute standard t-test significance (P-values shown) between NC-Cm ablated and Control delta Ct values. C) Schematic model in which NC-Cms provide spatial patterning of trabeculation, utilizing components of Notch/Neuregulin pathways in adjacent cardiomyocytes not derived from neural crest.

**Figure 4.**
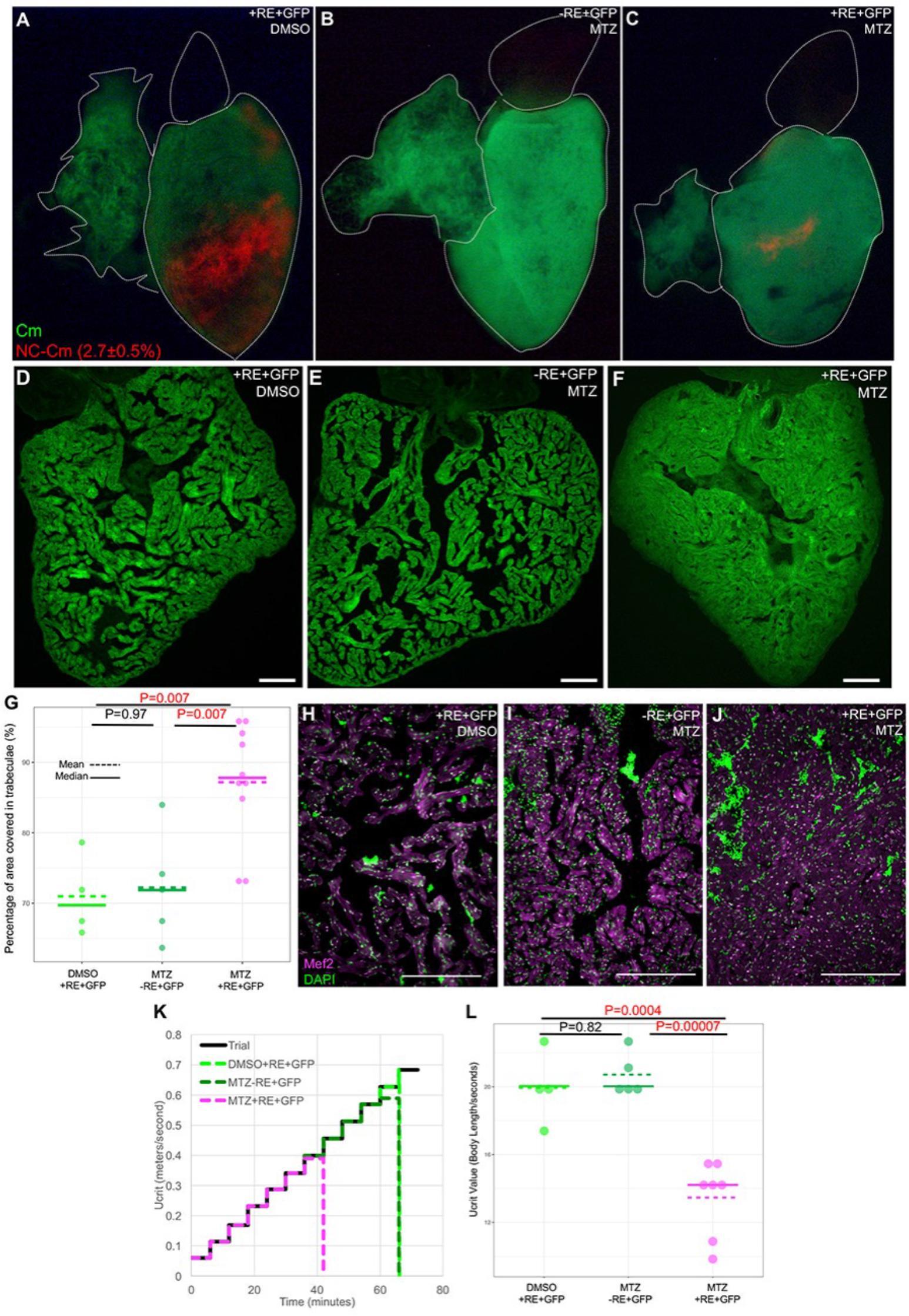
Ablation of NC-Cms during embryogenesis results in hypertrabeculated non-compaction cardiomyopathy and heart failure in adults. A-C) Whole mount fluorescent images of hearts isolated from adult zebrafish (4-month-old) generated from embryonic NC-Cms ablation experiments, as in Figure 2A. Numbers represent percent of total (GFP+) cardiomyocytes, not total heart cells, quantified by flow cytometry (FACS) of dissociated, individual +RE+GFP, untreated adult hearts (see Extended Figure 6). D-F) Fluorescent microscopy sections of hearts from sibling individuals as in A-C. scale bar = 100uM. G) Quantification of area of ventricle covered in trabeculae from sections similar to D-F. Dots represent sibling individuals from each condition pooled from biological replicates. Solid line = median of data, dashed line = mean. P values computed by TukeyHSD on ANOVA (F(2,16)=9.48). H-J) Microscopy sections of ventricles stained with Mef2 and DAPI. Nuclei that are both Mef2 and DAPI positive were used to count the number of cardiomyocytes per trabeculae area (see Extended Fig7A). Scale bar = 200uM. K) Swim trial assay with incremental speed increases of 0.05m/s every 6 minutes (solid black line). Average assay results for DMSO control (light green, n=4), MTZ control (dark green, n=5) and NC-CM ablated (magenta, n=8) adults. Sibling males were used as controls and data shown are amalgamated from multiple biological replicates. See Extended Movie 2. L) Individual Ucrit results normalized to body length from assay in H. Dots represent individual males from each condition. Solid line = median of data, dashed line = mean. P values computed by TukeyHSD on ANOVA (F(2,13)=24.65).

**Extended Figure 1.**
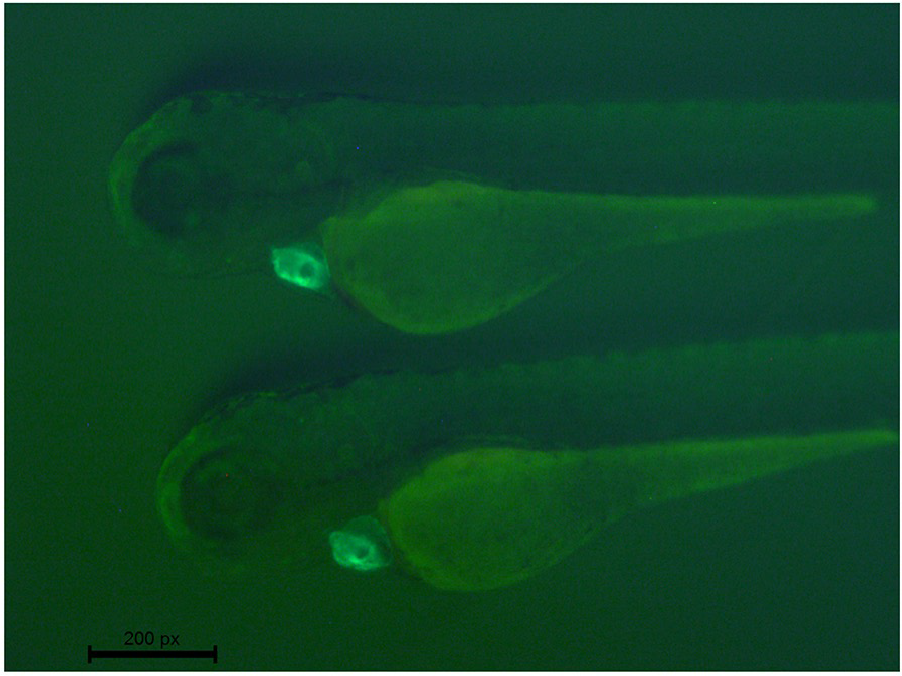
Whole mount fluorescent images of 4dpf *Cm:KillSwitch* transgenic embryos showing exclusive GFP fluorescence in the heart.

**Extended Figure 2.**
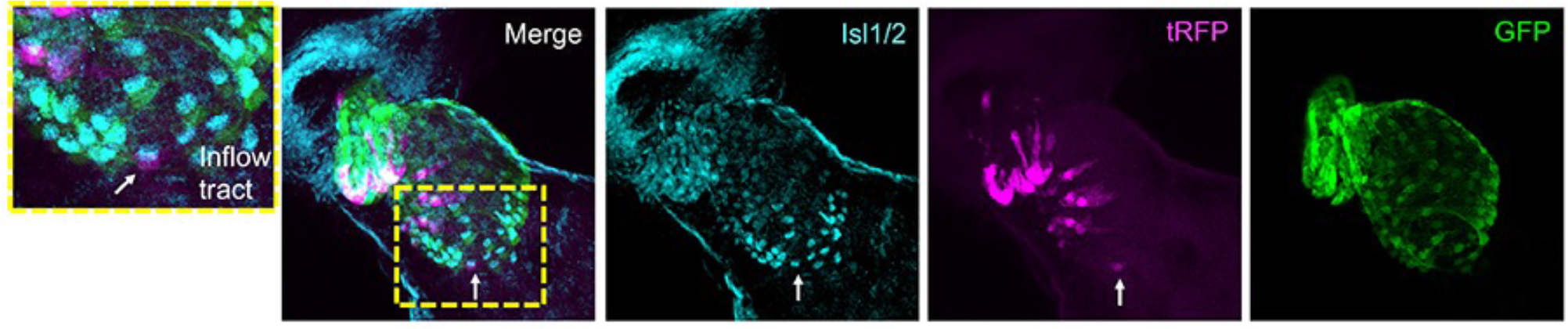
Isl1/2 antibody staining co-localizes with some NC-Cms indicating integration of neural crest cells into anterior second heart field. Arrow indicates single NC-Cm in the inflow tract that is positive for Isl1/2, a marker of pacemaker cells^27^. tagRFP and GFP were detected by immunolabeling.

**Extended Figure 3.**
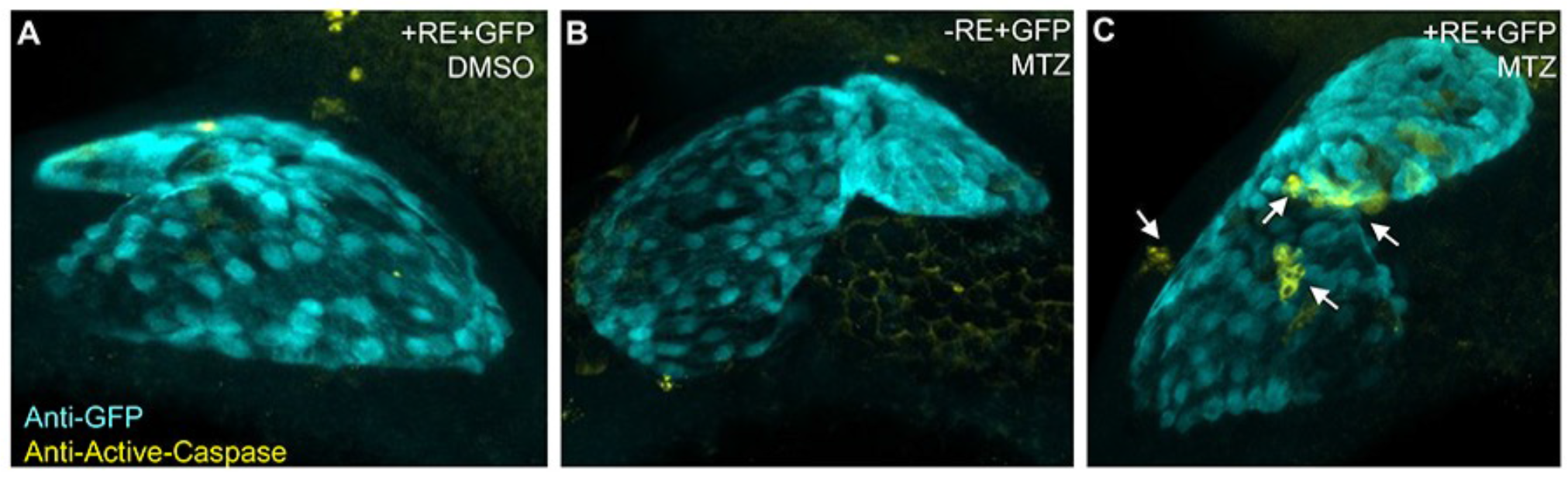
Cell death in NC-CMs after MTZ treatment. Embryos derived from transgenic crosses as described in Figure 1 and 2A. A) Control: DMSO treated double transgenic siblings. B) Control: MTZ treated single transgenic siblings. C) MTZ treated double transgenic (+RE+GFP) had positive staining for Active Caspase 3, indicative of cell death (arrows), that was not observed in hearts from controls.

**Extended Figure 4.**
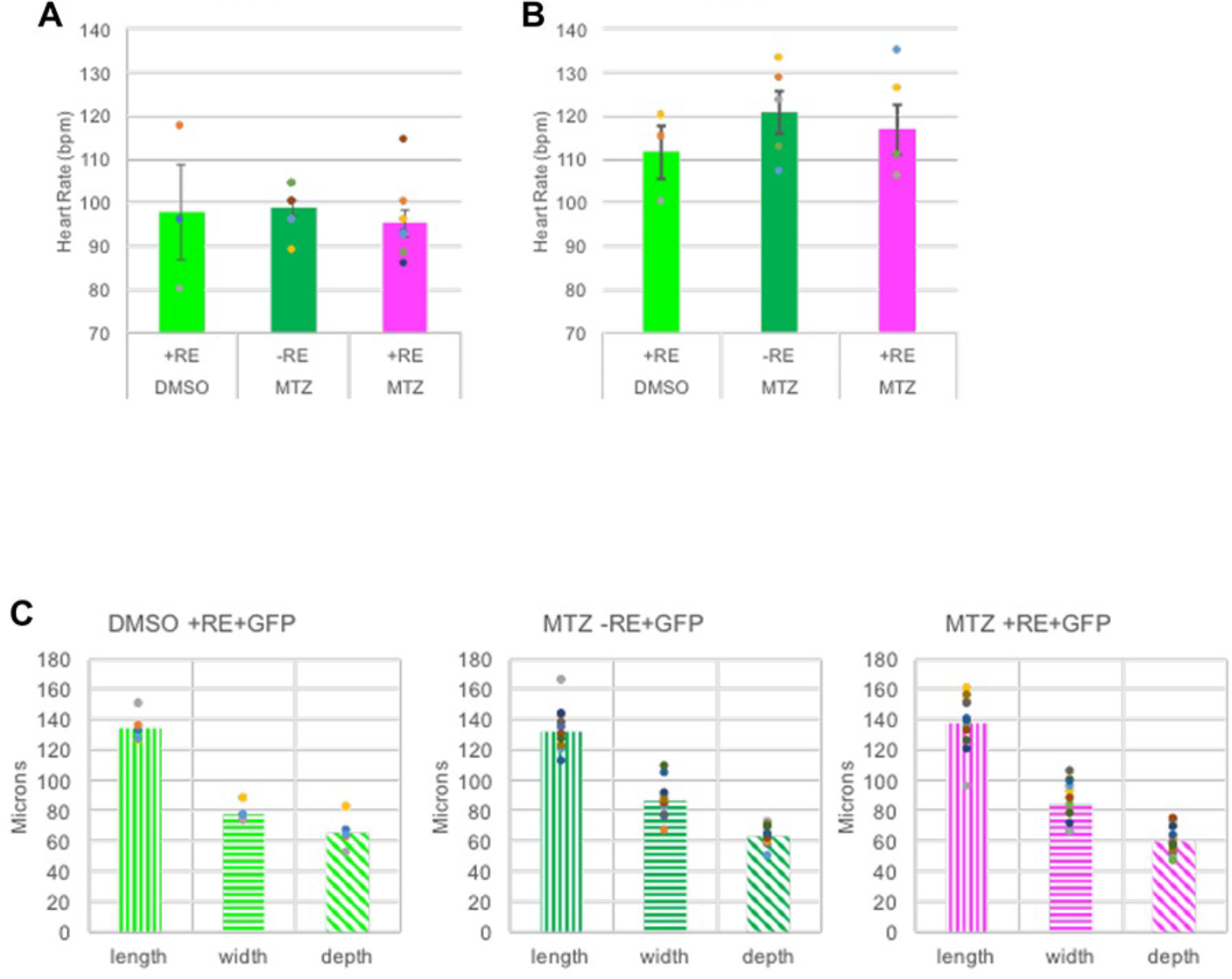
NC-Cm ablation has no significant effect on embryonic and juvenile heart rate or size. Heart rate (A) in 6dpf embryos and (B) in 14dpf juveniles. C) Ventricle dimension measurements from controls and MTZ treated siblings.

**Extended Figure 5.**
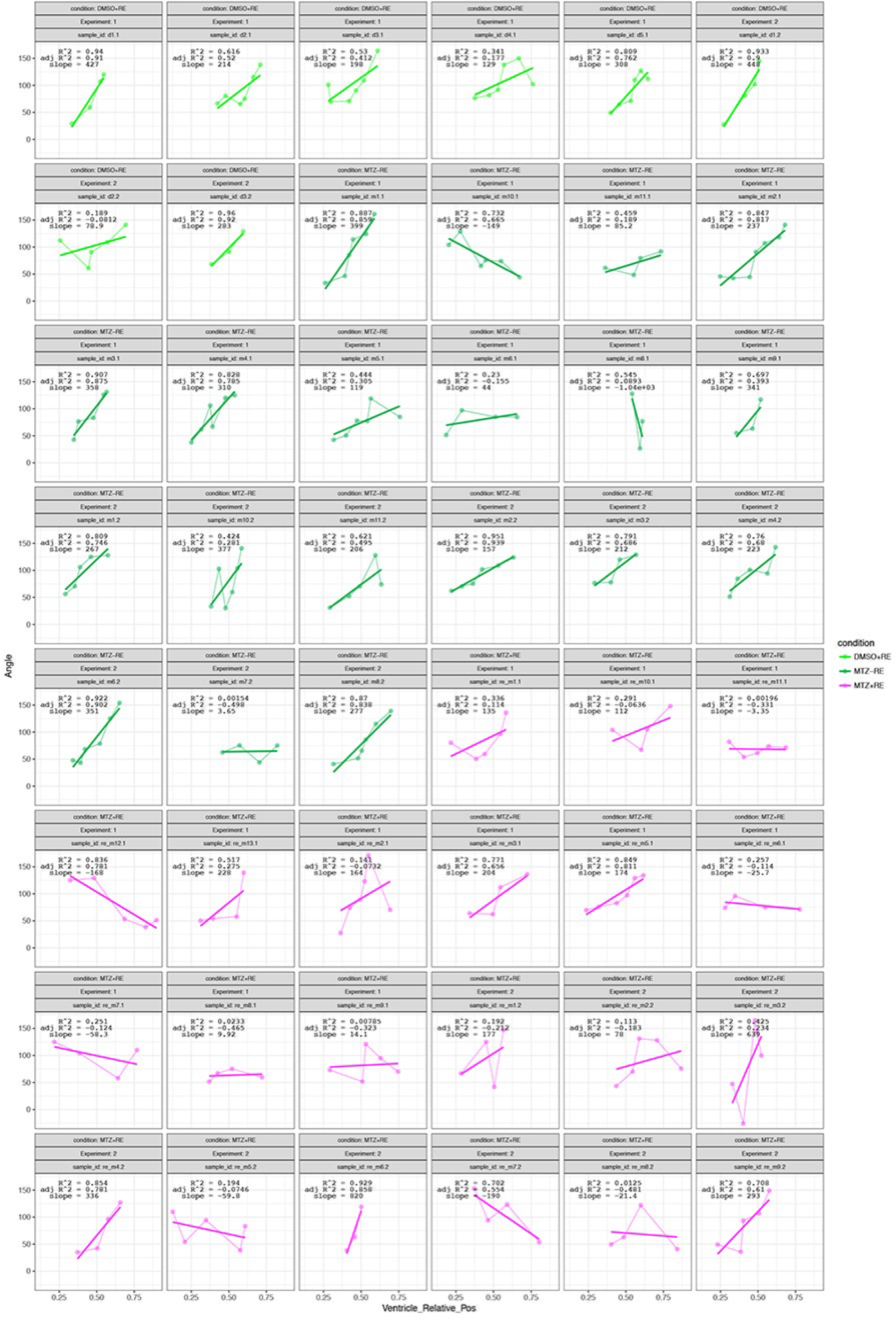
Slope calculations from angle and anterior-posterior position measurements of trabeculae in individual hearts from experiments described in Figure 2. Each plot is an individual heart. Light green is DMSO +RE+GFP, dark green is MTZ −RE+GFP and magenta is MTZ +RE+GFP conditions. Data shown are from one biological replicate. Similar quantification was done for two other biological replicates and shown in sum in Figure 2F.

**Extended Figure 6.**
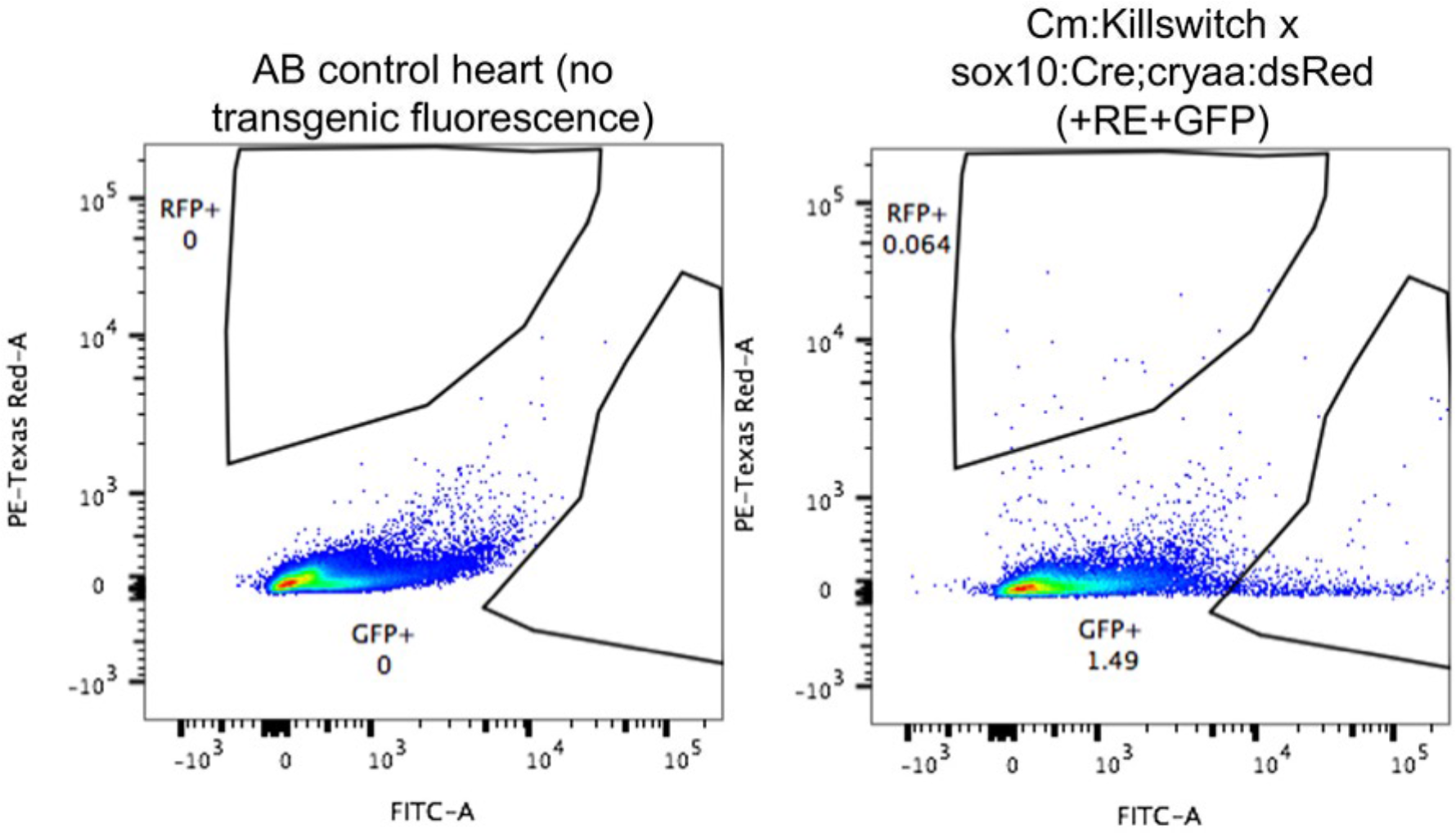
Flow cytometry analysis and quantification of labeled cardiomyocytes in *Cm:KillSwitch x Tg(sox10:cre;cryaa:dsRed)* individual adult heart versus AB control. GFP and RFP gates were drawn based on non-transgenic/non-fluorescent AB wild-type hearts. Dissociated cells were gated on viability (DAPI) and singlets before analyzing GFP and RFP populations. Three individual hearts were similarly analyzed by flow cytometry to quantify numbers of GFP+ and RFP+ cardiomyocytes. GFP+ and RFP+ percentages were added to quantify total number of cardiomyocytes in a whole heart and then RFP+ numbers were divided by this to generate their percent contribution to the total adult cardiomyocyte population. The average values of these quantifications are listed in Figure 4A. Heart dissociation protocol was carried out as previously described^28^.

**Extended Figure 7.**
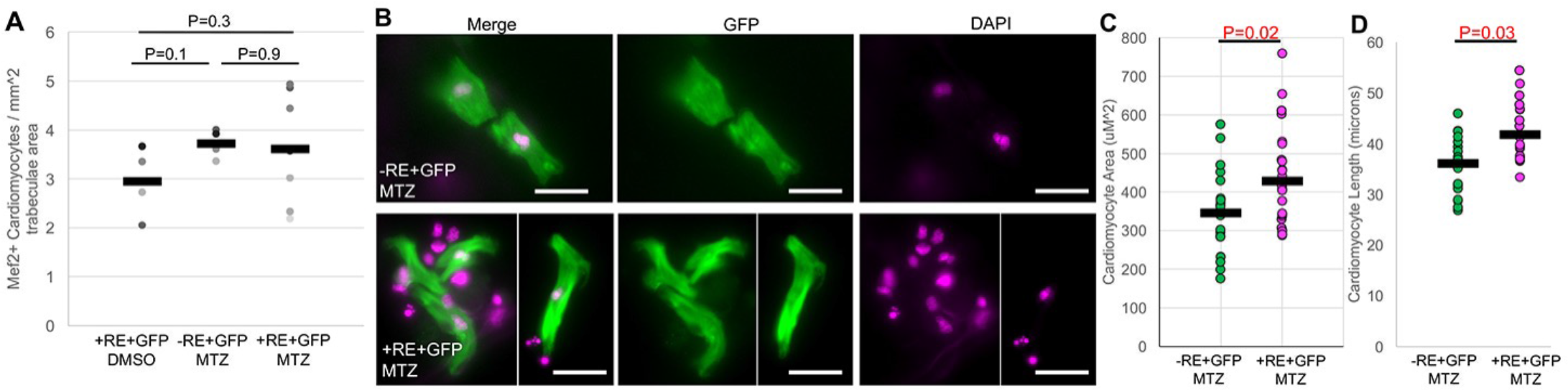
Quantification of cardiomyocyte number and size in adult hearts. A) Cardiomyocyte number per trabeculae area as quantified from microscopy section examples in Figure 4H-J. Dots represent individual adult section measurements and bars the mean of each sample. P-values on from two-sample standard T-tests. B) Examples of cardiomyocyte chamber cultures from NC-Cm ablated (+RE+GFP, MTZ, bottom panel) and sibling control (-RE+GFP, MTZ, top panel). GFP and DAPI staining of example cardiomyocytes from each sample. Note GFP negative, DAPI positive cells are present in images and represent non-cardiomyocyte cells of the dissociated ventricle cell population. Scale bar = 20uM. C) Quantification of individual cardiomyocyte area from chamber cultures as in ‘B.’N ≥ 25 cells per sample. P-value from standard t-test. D) Quantification of individual cardiomyocyte length from chamber cultures as in ‘B.’

**Extended Figure 8.**
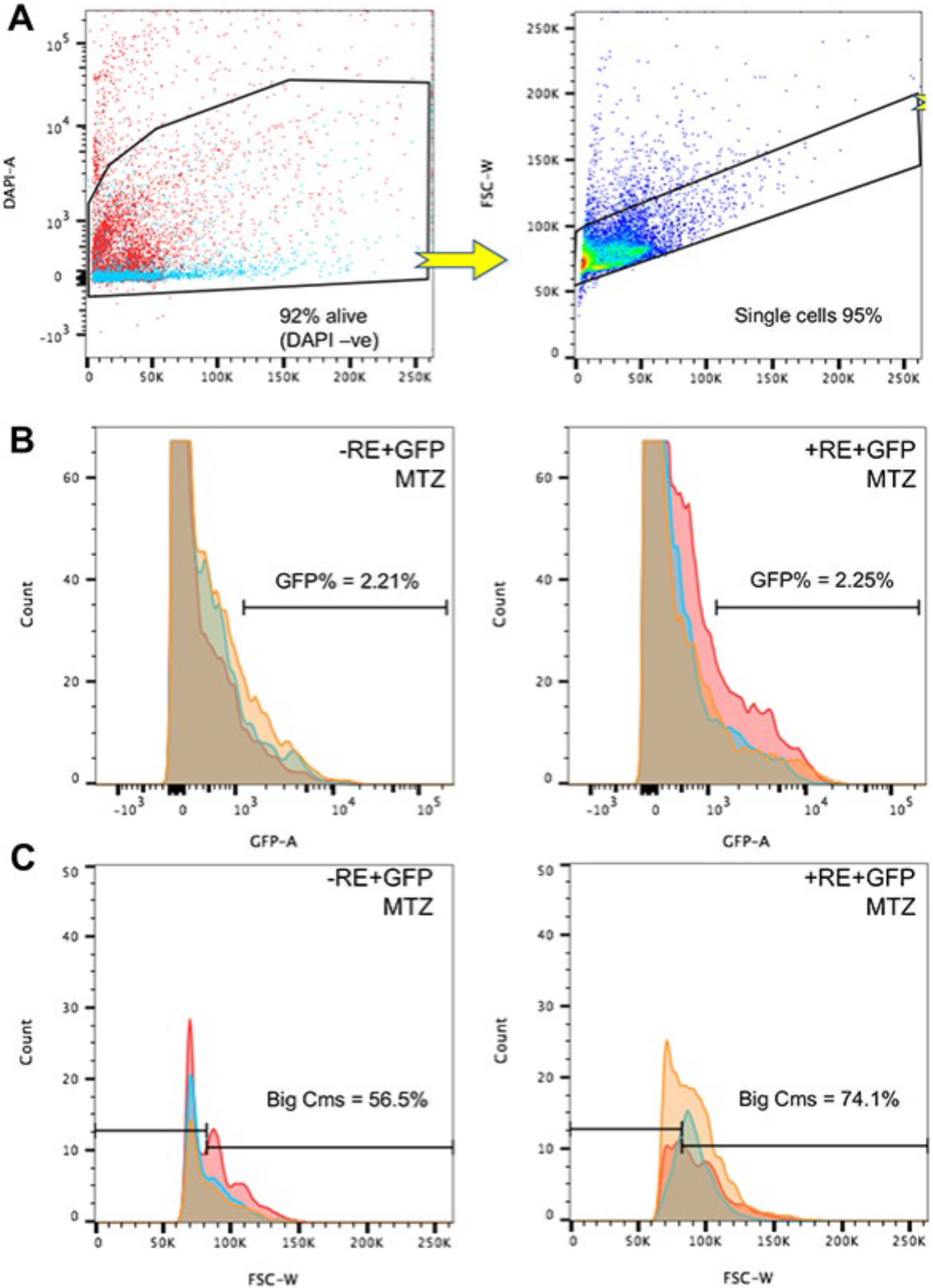
Flow cytometry analysis of NC-Cm ablated and sibling control dissociated ventricles. A) Flow cytometry gating and analysis strategy for dissociate ventricle cells. Cells were first gated based on viability using DAPI staining, followed by selection for single cells as opposed to doublets based on the forward scatter width profile. Cells colored in blue represent the negative control for gating analysis, except for in the single cell gate. Percentages are averages of three or more individual samples in the full experimental analysis. B) Cell populations from the single cell gate in panel A were analyzed for their percentage of GFP positive cells by histogram analysis and gating in the GFP channel, using the negative control sample to set the GFP+ gate. Percentages shown are averages from n ≥ 3 individual, dissociated ventricle analysis in each sample and were not significantly different (p=0.93, standard t-test, control = left and NC-Cm ablated = right). Colors in the histogram represent individuals in each sample analysis and overlaid onto a single plot. C) GFP positive cells as determined from gating analysis in panel B were then analyzed for their size spectrum using the forward scatter width channel. The number of cells in the larger forward scatter width profile was not as prominent in the NC-Cm ablated samples (right panel). Thus, a gate was created based on the control forward scatter width profile to quantify the amount of larger GFP positive cardiomyocytes (‘Big Cm’s’ gate). A significantly increased proportion of large GFP+ cardiomyocytes were found in the NC-Cm ablated (+RE+GFP, MTZ) ventricle samples compared to control (p=0.009, standard t-test, percentages shown are averages of n ≥ 3 in each sample, colors represent individuals).

##### Extended Movie 1

3D reconstruction of a 3dpf heart from a transgenic embryo of *sox10:cre;cryaa:dsRed* and *Cm:KillSwitch*. NC-Cms are labeled in magenta and all other CMs are labeled in green.

##### Extended Movie 2

Example of an adult fish undergoing the swim trial test and collapsing to fatigue.

## Methods

Zebrafish were housed in accordance with IACUC policies. The AB genetic background was used for all experiments and lines generated.

### Transgenic line generation

The p5-entry clone for the 7.2 kb *sox10* promoter is described^29^. The *p5E-sox10* clone was used with zebrafish codon optimized Cre, called *pME-iCre*, *p3E-cryaa:dsRed* and *pDestpA* to generate a final construct, via Gateway LR recombination technology: *sox10:iCre;cryaa:dsRed*. To generate *myl7:loxP-eGFP-loxP-tRFP-2A-mNTR* (called *Cm:KillSwitch*), the *p5E-myl7*, *pME-floxedGFP, p3E-tRFP-2A-mNTR* and *pDestpA* were recombined with Gateway LR recombination. *P3E-tRFP-2A-mNTR* was generated by PCR isolation of the 2A-mNTR sequence from the plasmid *p3E-YFP-2A-mNTR* (gift of J.Mumm lab)^30^, followed by a fusion PCR with the tagRFP coding sequence. Final constructs were sequence verified and injected at 25-30ng/ul with 30ng of Tol2 mRNA into single cell stage AB embryos^31^. At least three founders were screened, verified for similar expression patterns of each transgene and outcrossed to AB adults to propagate the line. Heterozygotes for each transgene were used in all experiments described.

### Transgenic line manipulation

To ablate NC-Cms and demonstrate effectiveness/specificity of the *Cm:KillSwitch* line, different doses/incubation times of metronidazole (MTZ, Sigma cat. No. 46461) treatment were tested. Treatment of *sox10:Cre;cryaa:dsRed x Cm:KillSwitch* embryos with 5mM from 30hpf-48hpf was comparable, in terms of yield of NC-Cm cell death, to 10mM MTZ treatment from 48-56hpf. For all reported experiments MTZ treatment was then carried out at 5mM doses from 30hpf-48hpf. MTZ stock was resuspended in DMSO at 1M concentration followed by dilution in E3 embryo media to achieve the correct dose (5 or 10mM). MTZ stock was stored at 4C in the dark but not used older than a week after resuspension. The MTZ treatment regimens tested resulted in increased, specific and detectable cell death immediately after treatment (Extended Figure 2).

### Flow cytometry

Adult ventricle or whole hearts were dissected out of anaesthesized fish and placed into cold HBSS + 1%FBS media. Hearts or ventricles were then allowed to pump for a few minutes in order to release any blood and squeezed gently with forceps to remove blood. For analyzing only ventricles, the atrium and outflow tract were manually dissected away with forceps. Otherwise whole hearts or ventricles were cut into pieces and placed into Liberase DH (1mg/ml) containing HBSS + 1% FBS solution. They were then placed in a 28C shaking incubator to dissociate for ~15-20mins. Pipetting every 5 minutes was used to aid in dissociation. ~200uL of dissociation solution was used for 1-3 hearts. Dissociated samples were spun down at 2500 rpm for 5 mins, supernatant removed without disturbing the cell pellet and then resuspended with 350ul of HBSS + 1% FBS, placed on ice, incubated with DAPI for a viability analysis, and processed for flow cytometry analysis on a BD FACS Canto.

### Immunofluorescence and Microscopy

For endogenous transgenic fluorescence detection and imaging, embryos were incubated in E3 media with PTU addition to prevent pigment formation. Embryos were briefly treated with 0.5M KCl to relax hearts and then immediately fixed in 2%PFA+PBS for 1 hour at room temperature. Embryos were washed and mounted in low melt agarose for microscopy. 3D images were acquired on a Zeiss LSM 880 under fast mode and 20X magnification.

Antibody staining for anti-active-caspase3 was done according to the published protocol^32^. Briefly Rabbit-anti-active-Caspase3 antibody was used (1:200, BD Pharmingen 559565) with Chicken anti-GFP (1:1000, Aves labs) in 4% PFA fixed embryos permeabilized with 100% cold methanol for 2 hours at − 20C. Washes used PBS + 3% Tritonx100.

Antibody staining for Isl1/2 in the NC-Cm labelled embryos was carried out as described^10^. Embryos were fixed in 2% PFA+PIPES buffer and incubated with Rabbit anti-tRFP (1:200, Life technologies R10367.) and Chicken anti-GFP.

Antibody staining for MF20 (DSHB) was carried out as described^33^.

Antibody staining for Mef2 (Abcam 64644) on ventricle sections was carried out on 4% PFA+PBS, fixed adult hearts that were cryosectioned into 10uM sections. Sections were boiled in citrate buffer for
~40mins, washed with PBS + 0.3% Triton x100 (PBT), blocked with PBT + 5% goat serum, 1% DMSO and 5ug/ml BSA. Antibody staining was carried out at 1:200 in blocking solution overnight at 4C. Washes used PBT and secondary antibody staining utilized AlexaFluor 568 goat anti-rabbit (catalog no.) in blocking solution.

Chamber cultures of dissociated adult ventricles were carried out by dissociating pools of isolated ventricles similar to the flow cytometry protocol above. The resuspension was then distributed into a single chamber of a Lab-Tek 8-chamber, chamber slide and incubated for 24hrs at 28C to allow settling and adherence of cells. To fix cultured cells for microscopy, media was largely removed but never left completely dry and 4% PFA +PBS was added carefully to not disturb adherent cells and incubated for 10mins. Chambers were washed in PBS and processed for staining with GFP antibody as in above methods.

### Image Analysis

Imaris software (v8.4.1) was used to reconstruct 3D microscopy images and count cell numbers. Imaris “surfaces” was used to analyze the volume of NC-Cm contribution by generating a surface for the entire RFP channel and assessing the volume compared to the volume of the GFP channel + RFP channel (combined they were considered the whole Cm volume of the heart). The “clipping plane” feature of Imaris was used to view individual trabeculae in embryonic ventricles and generate angle measurements using the “measurement points” feature. Only ventricles with greater than 3 measurable angles were used for analysis.

Trabeculae angles and relative heart position data were gathered for embryonic ventricles from Imaris analysis. These data were input into R programming interface to compute a slope value for each heart measured. Slopes were computed using the ‘lm’ function (linear regression model) in R, on individual ventricle measurements. Individual heart data for angle by position measurements are displayed in Extended figure 5 along with their computed slope and regression value.

Measurements of trabeculation in adult ventricle sections were generated using Fiji software. GFP fluorescent images (GFP fluorescence due to Cm or trabeculae structures of the ventricle) of adult ventricle sections were thresholded for a uniform value of intensity and then applied across all samples. A uniform area selection was used across all section samples to generate the percent of ventricle area covered in GFP/trabeculae.

Fiji was used to quantify the number of Mef2+, DAPI+ nuclei within the adult ventricle trabeculae. Trabeculae were outlined and selected manually to create an ROI and Mef2+, GFP+ nuclei were counted within each ROI, followed by a measurement of the ROI area. The number of nuclei were divided by the area in millimeters squared to tabulate values graphed in Extended Figure 7A.

For analysis of cardiomyocyte size in chamber cultures of dissociated ventricles, Fiji, was used to manually outline the GFP+ cardiomyocyte images acquired by compound fluorescent microscopy. DAPI was used to confirm a single cardiomyocyte was being analyzed and outlined for area measurements. To calculate length and width of cardiomyocytes the longest line was drawn from edge to edge length or width wise and then measured in micron units. Width measurements were not significant between NC-Cm ablated cardiomyocytes and sibling controls and area and length measurements are graphed in Extended Figure 7 C and D.

### Swim trials

A Loligo Systems swim tunnel setup was used to test individual adult males greater than 4 months of age for swim trial performance. Swim speeds in meters per second were based on calibrated instrument setting measurements. Incremental step sizes of 6 minutes and Ucrit values and measurements were calculated as described previously^34^. Fish were considered fatigued based on their inability to remove themselves from the mesh tunnel end for greater than 4 seconds. Body length normalization of Ucrit values was generated by measuring the length of the fish. Data shown are a compile of multiple biological replicates and sibling cohorts.

### Quantitative PCR

4dpf hearts were extracted from treated and control siblings from the *Cm:KillSwitch* cross to *Tg(sox10:Cre;cryaa:dsRed)* as described previously^35^. These hearts were immediately lysed in trizol and processed for RNA using a Zymo mini RNA kit. The total RNA was used for cDNA synthesis with BioRad iScript 5X master mix. Subsequent cDNA was used in multiplex qPCR reactions with the following gene primers: *rpl11, myl7, jag2B, erbb2, nrg2A*. Primer and probe sequences are listed in Extended data table 1. Delta Ct calculations were normalized to *rpl11* expression levels and control sibling expression levels.

### Statistics

R graphic programming was used to generate the dot plots in Figure 2F, 3G and 3I. ANOVA tests were run on dot plot data in 2F, 3G and 3I to test significance.

Student T-tests were performed on qPCR data in 3B on delta Ct values from control and NC-Cm ablated heart expression values.

**Extended data table 1.**
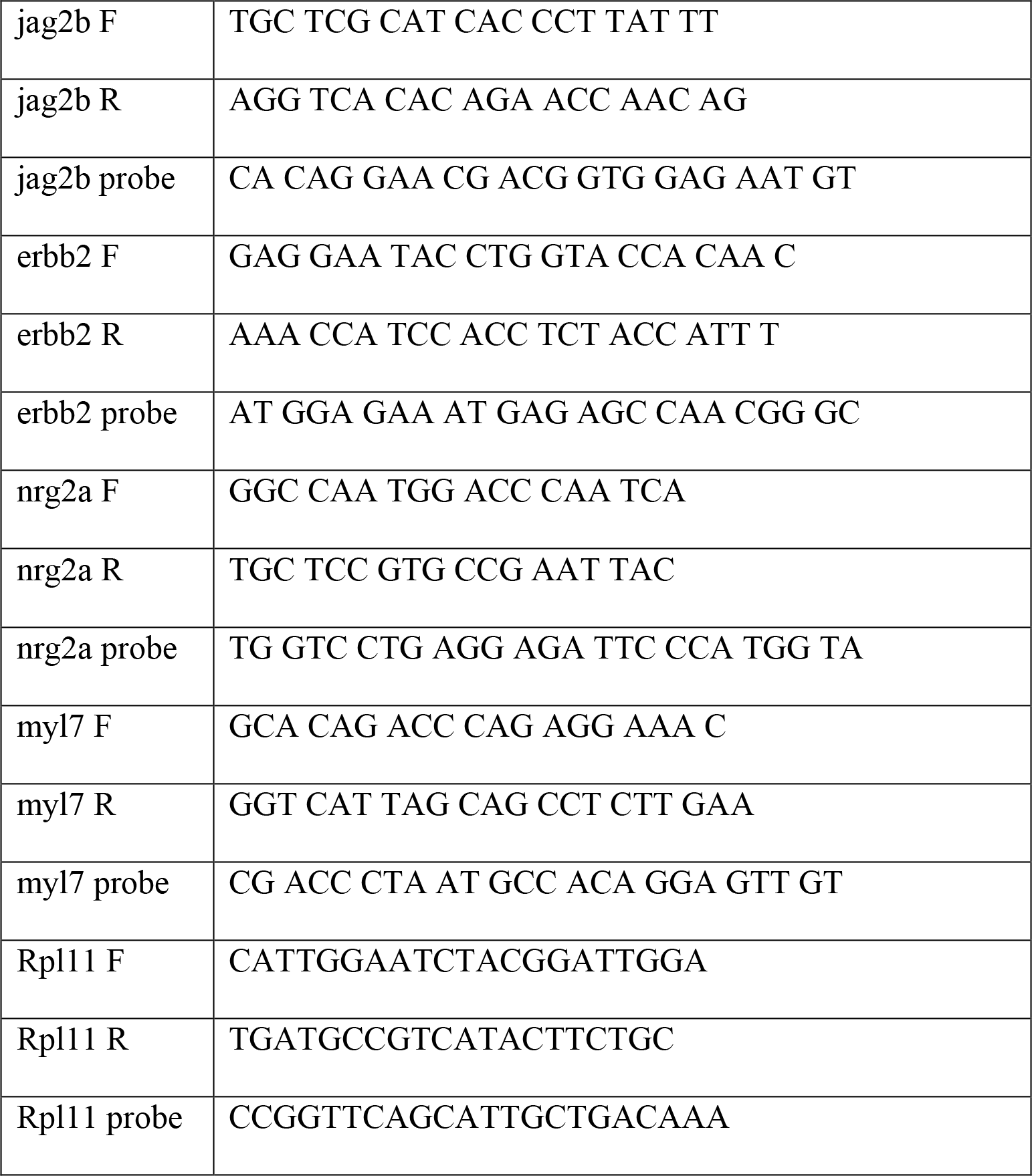
Primers and Probes used in qPCR.

## Acknowledgements

We gratefully acknowledge Dr. Jerald Johnson, BYU, for lending us a swim tunnel for these studies. We thank Dr. Jeffrey Mumm and Dr. Kristen Kwan for providing reagents. We thank Dr. Maureen Condic, Dr. Rodney Stewart, Dr. Marti Tristani for their review of this manuscript, and Yost lab members for discussions. This study was funded by a NHLBI Bench-to-Bassinet Consortium (http://www.benchtobassinet.com) grant to HJY (UM1HL098160). SAW was supported by National Institutes of Health under Ruth L. Kirschstein National Research Service Award 2T32HL007576-31 from the National Heart, Lung, and Blood Institute.

## Author contributions

SAW conceived, designed and conducted the experiments with input from HJY. HJY and SAW wrote the manuscript. BLD carried out statistical analysis and graphical presentation of data.

